# Acute Ketamine Modulated Functional Brain Coupling and Dissociative and Affective States in Human Subjects: Interim Analyses

**DOI:** 10.1101/2021.09.20.460992

**Authors:** Laura M. Hack, Katherine G. Warthen, Xue Zhang, Boris D. Heifets, Trisha Suppes, Peter van Roessel, Carolyn I. Rodriguez, Brian Knutson, Leanne M. Williams

## Abstract

Ketamine is a non-competitive antagonist of the N-methyl-D-aspartate (NMDA) glutamate receptor that is both a drug of abuse and an FDA-approved anesthetic used off-label for treatment-resistant depression. Despite its growing clinical use for depression and pain, the relationships between the acute dissociative and affective effects of ketamine that contribute to its abuse liability and therapeutic potential, along with the neural mechanisms underlying these effects, are not well established. To address this need, we have implemented a randomized, double-blinded, placebo-controlled, within-subjects mechanistic trial. Healthy adult subjects undergo infusion with two fixed doses of subanesthetic racemic intravenous (IV) ketamine and placebo and their acute responses are assessed with self-report questionnaires, behavioral measures, hormone levels, and neuroimaging. As planned in our analysis strategy, we present interim results for the first 7 subjects of our study, focusing on dissociative and affective states and resting functional brain coupling signatures of these states. The first key finding was that ketamine induced dose-dependent increases in dissociation and related intoxication. Ketamine also altered affective states, reducing emotional insensitivity but increasing stress assessed by cortisol. Second, ketamine had an effect on altering brain connectivity, particularly for specific connections between regions of the reward and negative affect circuits and involving thalamic sub-regions. Third, regarding brain-response associations, ketamine-induced increases in amygdala-anteroventral thalamus coupling were correlated with greater dissociation and intoxication, whereas decreases in the coupling of the anteromedial thalamus and posterior parietal thalamus were correlated with increased sensory aspects of reward responsiveness. Additional specific correlations were observed between affective measures relevant to reward responsiveness or its absence and drug-altered changes in localized functional connections involving the nucleus accumbens (NAcc), amygdala, and thalamic sub-regions. We also discovered a consistent profile of negative associations between ketamine altered connectivity involving the NAcc and specific thalamic sub-regions and effects of anxiety. Further, drug-altered increases in the coupling of the amygdala and anteroventral thalamus were associated with increases in cortisol, an indicator of biochemical stress. The findings highlight the utility of integrating self-reports, objective measures, and functional neuroimaging to disentangle the brain states underlying specific acute responses induced by ketamine. With the likely continued expansion of FDA indications for ketamine, understanding acute responses and underlying neural mechanisms is important for maximizing the therapeutic potential of ketamine while minimizing the risk of promoting misuse or abuse of this substance.

## Introduction

Ketamine is a non-competitive antagonist of the N-methyl-D-aspartate (NMDA) glutamate receptor and an FDA-approved dissociative anesthetic that was introduced into clinical practice in the US in the 1960’s as a novel agent that could produce effective sedation and analgesia while maintaining cardiorespiratory stability [1]. By the late 1970’s, ketamine had become a heavily abused substance throughout the world. By the turn of the 21^st^ century, subanesthetic doses of ketamine began to show promise as a breakthrough, fast acting therapy for treatment-resistant depression (TRD). Throughout the 2000’s, ketamine has increasingly been used as an off-label therapy for TRD, especially among suicidal patients [2], with its use rising further after FDA approval of intranasal esketamine in March 2019 for TRD in adults and in August 2020 for depressive symptoms in adults with major depressive disorder with acute suicidal ideation or behavior. Although the acute effects of ketamine have been correlated with both the abuse liability and treatment efficacy of ketamine, an in-depth characterization of the relationships between the dissociative and affective responses to acute subanesthetic ketamine administration in healthy subjects and the neural correlates of these responses has not been completed.

One of the key acute experiences that ketamine induces is dissociation, which is broadly defined as feelings of disconnection from one’s body and/or environment. In the context of ketamine, it may include a general sensory detachment, mind-body uncoupling, and feelings of floating in the absence of delirium [3, 4]. While challenging to obtain estimates of the prevalence of dissociation in abuse settings, this effect is known to commonly occur at doses therapeutic for TRD. For example, in short-term phase III trials of esketamine for those aged 18-64 years old, 27% of subjects (total n = 346) experienced dissociation [5]. Anecdotally, dissociation is noted as a reason for abusing ketamine, along with hallucinations and feelings of intoxication, which is a broad term that, for ketamine, can include mild euphoria that has been likened to alcohol intoxication as well as feelings of calmness and sedation. Some individuals experience ketamine-induced dissociation associated with positive affective states, whereas for others, including those who do not like the sense of loss of control, it induces negative affective states [6]. There is mixed evidence as to whether dissociation is necessary for ketamine’s antidepressant effect. For example, a meta-analysis showed it was not necessary [7], whereas another study demonstrated a correlation between reduction in depression scores and the depersonalization aspect of ketamine-induced dissociation in particular [8].

Regarding affective states, deconstructing these states may also be essential for understanding the potential therapeutic versus adverse effects, especially given that these effects tend to show opposing directions of change in clinical and healthy subjects. Therapeutic studies of subanesthetic IV ketamine in depressed patients show that, as early as 25-40 minutes after starting an infusion, reductions in overall depressive symptoms, are observed in some individuals [9–11]. Interestingly, a study in healthy subjects showed that ketamine significantly *increased* depressive symptoms as early as 40 minutes post-infusion [12]. Ketamine may induce stress-related effects of ketamine, reflected in self-reports of increased anxiety in both depressed people [13] and in healthy subjects [12]. These increases may be due to the biochemical impact of the drug. Indeed, subanesthetic doses of ketamine have been shown to increase the stress hormone cortisol at least two fold within an hour of administration [6, 14, 15].

In human resting functional MRI studies, ketamine has been found to produce a brain-wide increase in resting functional connectivity (FC) between thalamus and cortical association and limbic circuits [2, 16–18]. Graph theoretical analyses of resting brain states have revealed that ketamine induces a global functional reorganization of circuit connectivity, shifting from a distributed network of connections to a more subcortically focused connections [18]. At a local level, acute ketamine infusion has been found to induce a decrease in resting-state FC between the locus coeruleus and the thalamus with peak decreases at particular subthalamic nuclei [19]. Ketamine administration has also been associated with changes in the FC of the hippocampus in both mice and humans [20, 21].

Although substantial progress has been made in understanding the acute effects of ketamine we lack data on how to disentangle the dissociative from affective effects of the drug, and whether these effects are supported by distinct alterations in brain states. For future treatment applications it will be important to have a fine-grained understanding of which dissociative and affective responses to ketamine are therapeutic and which may be aversive and the extent to which these responses are personalized and vary between individuals. Because the findings to date indicate that dissociation may be a potentially positive experience in some instances, associated with a therapeutic outcome, whereas in others it may be an indicator of adverse experiences, parsing these effects at an individual level is essential.

Our focus on humans is guided by a translational neuroscience approach. We are informed by findings from rodent models and, in reverse, seek to translate our human findings back to these models. With respect to dissociative experiences, in seminal work, Deisseroth and colleagues observed that ketamine induced rhythmic activity in the retrosplenial cortex (RSP) of rodents indicative of dissociation, which correlated with activity in the laterodorsal and anteroventral thalamus but decorrelated with activity in the anteromedial thalamus and somatosensory cortex [22]. Human neuroimaging trials suggest that dissociative experiences may also involve local changes in correlated medial prefrontal cortical (mPFC)-striatal activity [23], which specifically utilize glutamatergic transmission [24]. The nucleus accumbens (NAcc) FC with the ventromedial prefrontal cortex has been reported to increase in a group of healthy subjects given ketamine [23].

Thus, our overall goal is to provide an in-depth characterization of dissociative and affective responses to acute (during and within minutes of the infusion) ketamine in healthy subjects, and the concurrent altered functional brain connectivity signatures of these acute responses. To achieve this, we conducted a double-blinded placebo-controlled human mechanistic trial by utilizing two doses of IV ketamine and saline placebo infused over 40 minutes: a higher dose (0.5mg/kg), which is standardly used in therapeutic settings for the treatment of TRD, and a lower dose (0.05mg/kg) for comparison purposes.

Here, we report the preliminary results from this study in 7 out of a planned final sample of 20 healthy subjects. These results derive from a pre-specified analytic strategy outlined in our protocol and when registering the study (NCT03475277– clinicaltrials.gov). Specifically, during real-time administration of two different doses of racemic IV ketamine, we aimed first to quantify ketamine-induced altered states involving dissociative experiences and affective responses. We assessed affective responses within domains of reward responsivity, negative affect and stress, using both self-reports and cortisol sampling. Second, we used high temporal resolution resting-state functional magnetic resonance imaging to quantify ketamine-induced altered brain states involving changes in resting-state FC. We then quantified brain-wide changes in global FC as well as localized changes in specific functional connections involving regions of reward circuitry and negative affect circuitry. Motivated by the seminal finding that specific profiles of coupling and uncoupling involving sub-regions of the thalamus characterize dissociative states in rodent models [22], we also focused more specifically on connections involving the thalamus and its subregions. Third, we quantified associations between subjects’ ketamine-altered dissociative and affective states and the underlying alterations in functional brain states. Analyses addressing these objectives focused on group average data as well as the signatures for each individual subject contributing to the average.

## Methods

### Participants

As planned, interim analyses focus on 7 participants between the ages of 18 and 55 years old who reported > 2 prior uses of ketamine. For a full list of eligibility criteria, see Table 1.

**Table 1.**
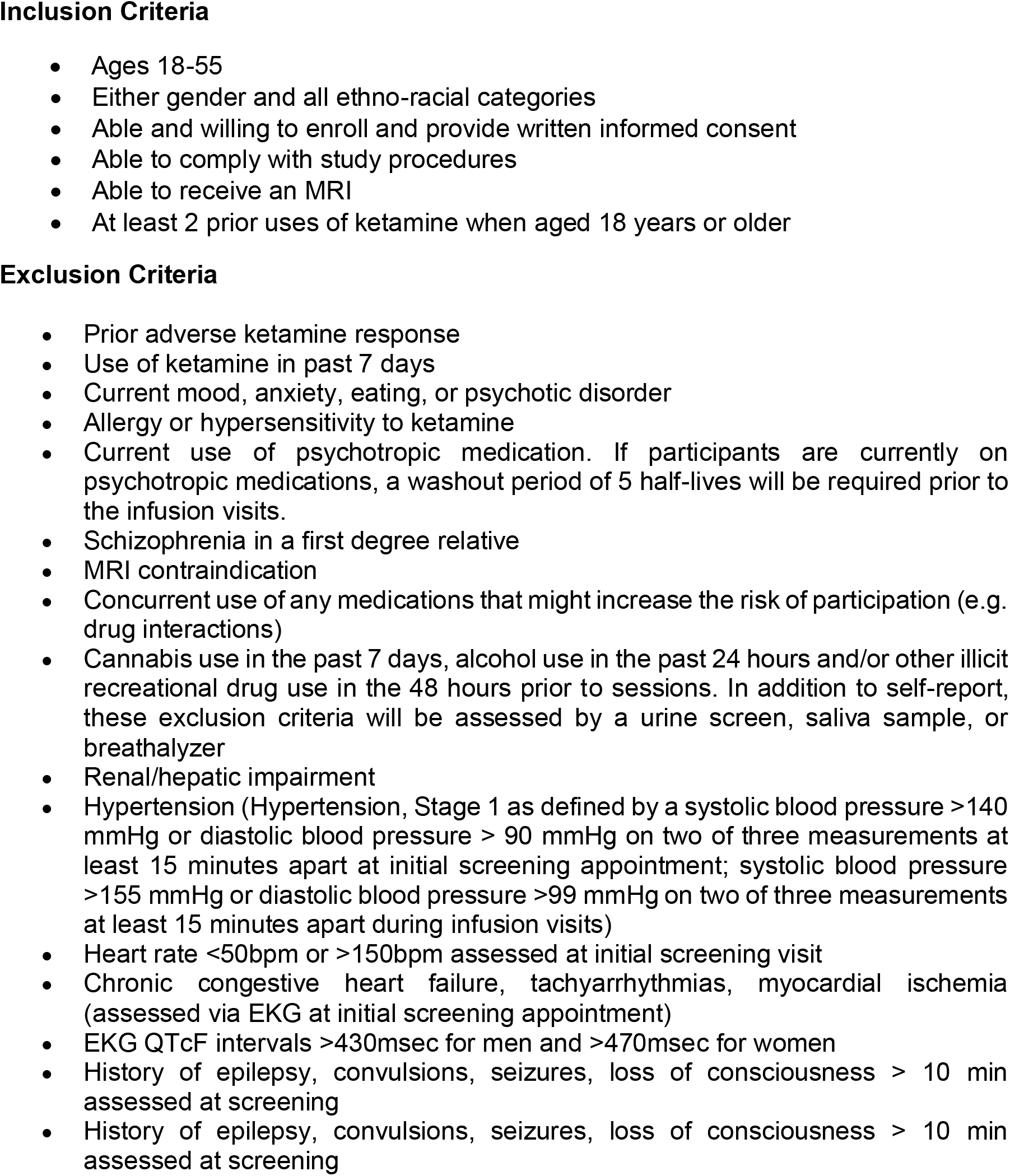
Inclusion and exclusion criteria

All participants were healthy adults with minimal psychiatric symptoms at baseline. Demographic characteristics and level of clinical anxiety and depressive symptoms are summarized in Table 2. Participants were recruited through Facebook Ads using IRB-approved material. Individuals who expressed interest in the study were directed to an online screening survey in REDCap and individuals who were eligible to participate were contacted by a research coordinator for a phone screening. On this phone call, research coordinators provided the individuals with additional information about the study, collected additional demographic information, and scheduled an in-person screening visit at a research clinic. Screening procedures were conducted by trained medical professionals including phlebotomists (blood draw), research nurses (ECG and vital signs), trained research coordinators (drug and psychiatric histories), and licensed study physicians (physical examinations and confirmation of psychiatric histories). Baseline and infusion session procedures were conducted by trained research coordinators (study assessments), research nurses (peripheral IV catheter placement), and licensed study physicians (ketamine infusions, monitoring, and safety assessments). Informed consent procedures were conducted by a trained research coordinator at the start of in-person screening visits. Travel was arranged for participants to all infusion visits and they were financially compensated for their participation in the study.

**Table 2.** Demographics and baseline clinical depression and anxiety levels

|  | **Overall**<br>**(N=7)** |
| --- | --- |
| **Age (years)** |  |
| Mean (SD) | 28.1 (6.41) |
| Median [Min, Max] | 27.0 [22.0, 36.0] |
| **Gender** |  |
| Male | 4 (57.1%) |
| Female | 3 (42.9%) |
| **Hamilton Anxiety Scale** |  |
| Mean (SD) | 0.429 (0.535) |
| Median [Min, Max] | 0 [0, 1.00] |
| **Hamilton Depression Scale** |  |
| Mean (SD) | 0.429 (0.535) |
| Median [Min, Max] | 0 [0, 1.00] |
| **9-item Patient Health Questionnaire** |  |
| Mean (SD) | 1.57 (1.51) |
| Median [Min, Max] | 1.00 [0, 4.00] |
| **7-Item Generalized Anxiety Disorder Scale Score** |  |
| Mean (SD) | 1.14 (1.07) |
| Median [Min, Max] | 1.00 [0, 3.00] |

### Visits

Participants underwent a total of 5 visits, including the screening visit, baseline visit, and 3 infusion visits at which the participants, research coordinator, and study physician were blinded to the intervention. At any of the 3 infusion visits, participants received saline, 0.05mg/kg ketamine, or 0.5mg/kg ketamine in a randomized order. We use IV ketamine administration as it provides the most predictable dosing with 100% bioavailability. Given the within-subjects design of the study, each participant received all three of the specified doses across the duration of the trial. The three infusion visits were separated by 10-14 days to avoid drug carry-over effects. Participants arrived fasted for infusion visits in the morning to reduce the risk of emesis, and they completed a urine drug screen and pregnancy test (if applicable), had their baseline vitals recorded, and had a peripheral IV catheter placed at the Clinical Trial Research Unit (CTRU) at Stanford University. Participants were then escorted to the Stanford Center for Cognitive and Neurobiological Imaging (CNI), where they were met by the study physician and subsequently began the infusion. Racemic ketamine (0.05 mg/kg or 0.5 mg/kg) or 0.9% normal saline (Mariner Advanced Pharmacy Corp, San Mateo, CA) was delivered via an elastomeric pump (Braun Easypump^®^ ST/LT) over 40 minutes. Before, during, and after the infusion, blinded self-report and clinician-administered assessments were completed. Participants were then guided to the MRI suite to begin scanning at approximately 60 minutes after initiation of infusion. In all conditions, pulse, blood pressure, and pulse-oximetry were continuously monitored during the infusions and scanning sessions. Saliva samples were collected at five timepoints throughout the visit. After the 1.5 hour scanning session (including tasks not described in the present report), the participant completed self-report assessments. The total visit time for infusion visits was 6-8 hours. The overall timeline for a visit is shown in Figure 1.

**Figure 1.**
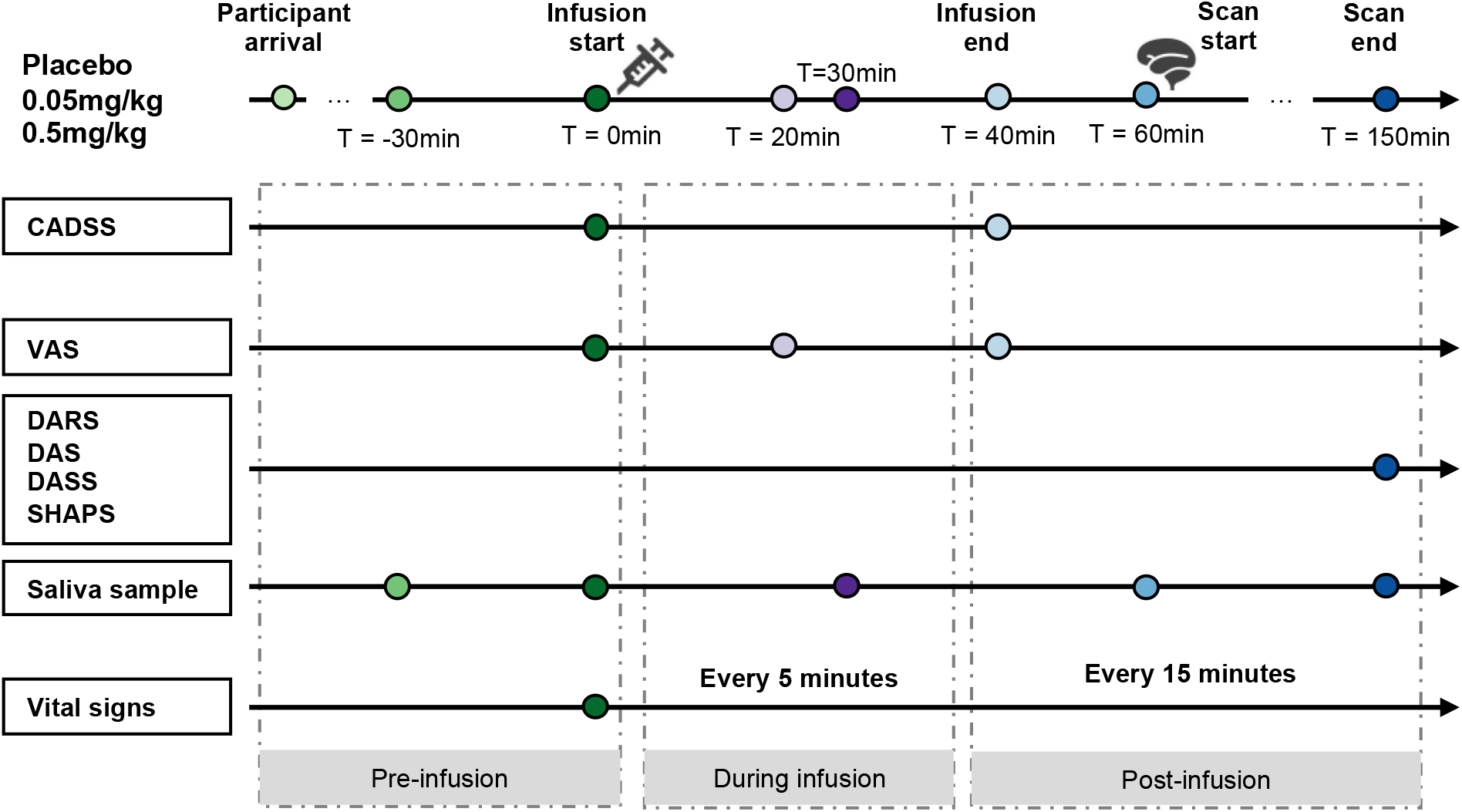
Study timeline. For each infusion visit, participants arrived in the morning, and they completed a urine drug screen and pregnancy test (if applicable), had their baseline vitals recorded, and had a peripheral IV catheter placed. Participants were then escorted to the scanning facility where they were met by the study physician and subsequently began the 40-minute infusion. After this, participants were escorted to the MRI suite to begin scanning. At multiple time points, the CADSS and VAS were used to assess dissociative states and related intoxication levels, the DARS, DAS, DASS (Depression and Anxiety scales), and SHAPS were used to assess affective states (related to reward responsiveness and negative affect), and the saliva sample was taken to assess affective states related to stress, along with the DASS Stress scale. Abbreviations: CADSS = Clinician-Administered Dissociative States Scale; VAS = Visual Analogue Scale for Intoxication; DARS = Dimensional Anhedonia Rating Scale; DAS = Dimensional Apathy Scale; DASS = Depression Anxiety Stress Scales; SHAPS = Snaith-Hamilton Pleasure Scale.

### Assessments of Dissociative and Affective States

We selected each of our measures to assess specific aspects of ketamine-altered dissociative and affective states, and we organize them according to the constructs as follows:

#### Dissociative States and Intoxication

##### Clinician-Administered Dissociative States Scale (CADSS)

A 23-item self-report questionnaire administered by a study clinician to assess present-state dissociative symptoms [25].

##### Visual Analogue Scale (VAS) for Intoxication

A single-item self-report question administered verbally by study staff asking a participant to rate their level of intoxication on a scale from 0-100 [26].

#### Affective Reward Responsiveness

##### Dimensional Apathy Scale (DAS)

A 24-item self-report questionnaire designed to assess dimensions of emotional apathy (extent of sensitivity through insensitivity), capacity for executive function (executive), and capacity for initiation of thoughts and behaviors (initiation) [27].

##### Snaith-Hamilton Pleasure Scale (SHAPS)

A 14-item self-report questionnaire designed to measure the dimension of pleasure through to loss of pleasure (anhedonia) [28]. This measure was scored on a 0-14 scale using the original scoring guidelines.

##### Dimensional Anhedonia Rating Scale (DARS)

A 17-item self-report questionnaire that examines domains of social interaction, interest in hobbies, interest in food and sensory states, with each item measured on a 5-point scale of 0 to 4. A high score is reflective of less anhedonia [29].

#### Negative Affect Mood and Anxiety States

##### Depression Anxiety Stress Scales (DASS, 42 item version)

A 42-item self-report questionnaire that assesses a continuum of experiences of depression, anxiety, and stress that occur in the population from minimal through moderate/severe using three subscales [30]. The DASS has been normed for use in community and patient groups and validated against other measures of anxiety such as the Beck Anxiety Inventory.

#### Negative Affect - Stress

##### DASS Stress subscale

This subscale was used to assess self-reported experiences of tension and stress [30].

##### Cortisol

Cortisol changes derived from saliva samples were used as a metric for quantifying biochemical stress altered by ketamine. Saliva samples were collected at five points during infusion visits (30 minutes before infusion, infusion start, and 30, 60, and 150 minutes after infusion) to measure cortisol levels. These samples were taken by placing a cotton swab into the participant’s mouth and allowing the swab to passively absorb saliva for 1-2 minutes, at which time the swab was removed and stored tubes in a securely locked laboratory facility. Samples were processed at the Dresden University of Technology in Dresden, Germany by liquid chromatography with tandem mass spectrometry.

### Brain Imaging

#### Resting-state Functional MRI

Participants were asked to look at a white crosshair on a black screen while allowing their mind to wander naturally for a five-minute session (posterior-anterior phase encoding direction), followed by a second five-minute session (anterior-posterior phase encoding direction). We employed a state-of-the-field multi-band approach based on Connectome quality imaging and which was implemented for neuroimaging data acquired at rest. These data were acquired on a 3T GE UHP MR scanner with a Nova Medical 32-channel head coil. We employed a multi-band acceleration of 6; acquisition time = 5:12, TE = 30 ms, TR = 0.71 s, FA = 54°, field of view(FOV) = 216 × 216 mm, 3D matrix size = 90 × 90 × 60, slice orientation = axial, angulation to AC-PC line, phase encoding = PA and AP, receiver bandwidth = 250 kHz, readout duration = 48.06 ms, echo spacing = 0.54 ms, number of volumes of resting-state =428, calibration volumes = 2, voxel size = 2.4 mm isotropic.

#### Structural MRI

A T1-weighted anatomical MRI image was collected for the purpose of normalization using the MPRAGE sequence. The parameters for the anatomical T1 image were as follows: TE = 3.828 ms, TR = 3s, FA = 8°, acquisition time = 8:33, field of view = 256 × 256 mm, 3D matrix size = 320 × 320 × 230, slice orientation = sagittal, angulation to AC-PC line, receiver bandwidth = 31.25 kHz, fat suppression = no, motion correction = PROMO, voxel size = 0.8 mm isotropic.

#### Image Preprocessing

Results included in this paper come from preprocessing performed using fMRIPrep 20.2.3 ([31, 32]; RRID:SCR_016216), which is based on Nipype 1.6.1 [33, 34]; RRID:SCR_002502).

#### Anatomical Data Preprocessing

A total of 1 T1-weighted (T1w) images were found within the input BIDS dataset. The T1-weighted (T1w) image was corrected for intensity non-uniformity (INU) with N4BiasFieldCorrection [35], distributed with ANTs 2.3.3 ([36], RRID:SCR_004757), and used as T1w-reference throughout the workflow. The T1w-reference was then skull-stripped with a Nipype implementation of the antsBrainExtraction.sh workflow (from ANTs), using OASIS30ANTs as target template. Brain tissue segmentation of cerebrospinal fluid (CSF), white-matter (WM) and gray-matter (GM) was performed on the brain-extracted T1w using fast (FSL 5.0.9, RRID:SCR_002823, [37]). Brain surfaces were reconstructed using recon-all (FreeSurfer 6.0.1, RRID:SCR_001847, [38]) and the brain mask estimated previously was refined with a custom variation of the method to reconcile ANTs-derived and FreeSurfer-derived segmentations of the cortical gray-matter of Mindboggle (RRID:SCR_002438, [39]). Volume-based spatial normalization to two standard spaces (MNI152NLin6Asym, MNI152NLin2009cAsym) was performed through nonlinear registration with antsRegistration (ANTs 2.3.3), using brain-extracted versions of both T1w reference and the T1w template. The following templates were selected for spatial normalization: FSL’s MNI ICBM 152 non-linear 6th Generation Asymmetric Average Brain Stereotaxic Registration Model [[40], RRID:SCR_002823; TemplateFlow ID: MNI152NLin6Asym], ICBM 152 Nonlinear Asymmetrical template version 2009c [[41], RRID:SCR_008796; TemplateFlow ID: MNI152NLin2009cAsym].

#### Functional Data Preprocessing

For each of the BOLD runs found per subject (across all tasks and sessions), the following preprocessing was performed. First, a reference volume and its skull-stripped version were generated by aligning and averaging 1 single-band references (SBRefs). Susceptibility distortion correction (SDC) was omitted. The BOLD reference was then co-registered to the T1w reference using bbregister (FreeSurfer) which implements boundary-based registration [42]. Co-registration was configured with six degrees of freedom. Head-motion parameters with respect to the BOLD reference (transformation matrices, and six corresponding rotation and translation parameters) are estimated before any spatiotemporal filtering using mcflirt (FSL 5.0.9, [43]). First, a reference volume and its skull-stripped version were generated using a custom methodology of fMRIPrep. The BOLD time-series were resampled onto the following surfaces (FreeSurfer reconstruction nomenclature): fsnative, fsaverage. The BOLD time-series (including slice-timing correction when applied) were resampled onto their original, native space by applying the transforms to correct for head-motion. These resampled BOLD time-series will be referred to as preprocessed BOLD in original space, or just preprocessed BOLD. The BOLD time-series were resampled into several standard spaces, correspondingly generating the following spatially-normalized, preprocessed BOLD runs: MNI152NLin6Asym, MNI152NLin2009cAsym. First, a reference volume and its skullstripped version were generated using a custom methodology of fMRIPrep. Grayordinates files [44] containing 91k samples were also generated using the highest-resolution fsaverage as intermediate standardized surface space. Automatic removal of motion artifacts using independent component analysis (ICA-AROMA, [45]) was performed on the preprocessed BOLD on MNI space time-series after removal of non-steady state volumes and spatial smoothing with an isotropic, Gaussian kernel of 6mm FWHM (full-width half-maximum). Corresponding “non-aggressively” denoised runs were produced after such smoothing. Additionally, the “aggressive” noise-regressors were collected and placed in the corresponding confounds file. Several confounding time-series were calculated based on the preprocessed BOLD: framewise displacement (FD), DVARS and three region-wise global signals. FD was computed using two formulations following Power (absolute sum of relative motions, [46]) and Jenkinson (relative root mean square displacement between affines, [43]). FD and DVARS are calculated for each functional run, both using their implementations in Nipype (following the definitions by [46]). The three global signals are extracted within the CSF, the WM, and the whole-brain masks. Additionally, a set of physiological regressors were extracted to allow for component-based noise correction (CompCor, [47]). Principal components are estimated after high-pass filtering the preprocessed BOLD time-series (using a discrete cosine filter with 128s cut-off) for the two CompCor variants: temporal (tCompCor) and anatomical (aCompCor). tCompCor components are then calculated from the top 2% variable voxels within the brain mask. For aCompCor, three probabilistic masks (CSF, WM and combined CSF+WM) are generated in anatomical space. The implementation differs from that of Behzadi et al. in that instead of eroding the masks by 2 pixels on BOLD space, the aCompCor masks are subtracted a mask of pixels that likely contain a volume fraction of GM. This mask is obtained by dilating a GM mask extracted from the FreeSurfer’s aseg segmentation, and it ensures components are not extracted from voxels containing a minimal fraction of GM. Finally, these masks are resampled into BOLD space and binarized by thresholding at 0.99 (as in the original implementation). Components are also calculated separately within the WM and CSF masks. For each CompCor decomposition, the k components with the largest singular values are retained, such that the retained components’ time series are sufficient to explain 50 percent of variance across the nuisance mask (CSF, WM, combined, or temporal). The remaining components are dropped from consideration.

The head-motion estimates calculated in the correction step were also placed within the corresponding confounds file. The confound time series derived from head motion estimates and global signals were expanded with the inclusion of temporal derivatives and quadratic terms for each [48]. Frames that exceeded a threshold of 0.5 mm FD or 1.5 standardised DVARS were annotated as motion outliers. All resamplings can be performed with a single interpolation step by composing all the pertinent transformations (i.e. head-motion transform matrices, susceptibility distortion correction when available, and co-registrations to anatomical and output spaces). Gridded (volumetric) resamplings were performed using antsApplyTransforms (ANTs), configured with Lanczos interpolation to minimize the smoothing effects of other kernels [49]. Non-gridded (surface) resamplings were performed using mri_vol2surf (FreeSurfer).

Many internal operations of fMRIPrep use Nilearn 0.6.2 ([50], RRID:SCR_001362), mostly within the functional processing workflow. For more details of the pipeline, see the section corresponding to workflows in fMRIPrep’s documentation.

##### Copyright Waiver

The above boilerplate text was automatically generated by fMRIPrep with the express intention that users should copy and paste this text into their manuscripts unchanged. It is released under the CC0 license.

#### Resting State Functional Connectivity Calculation

Resting state imaging data was processed with a custom toolbox in MATLAB [51, 52]. All included subjects had mean frame displacement <0.20 mm, and motion outliers detected using fMRIPrep were not above 25% of timepoints. Resting state data was parcellated into the regions of the Brainnetome (BNT) atlas (http://www.brainnetome.org/). The time series data from each region of interest (ROI) was quadratically detrended and COMPCOR and motion regression were performed using the confound regressors derived from fMRIPrep. Bandpass filtering was performed with a fast Fourier transform filter between 0.008 and 0.10 Hz. Pearson correlation coefficients were calculated between the average BOLD signals for each pair of brain regions and transformed into z-scores. Global efficiency was calculated with the BRAPH graph theory toolbox [53].

#### Predefined ROIs

Based on the pre-specified primary focus of our analysis strategy, informed by ketamine-induced changes in brain activity from preclinical and clinical literature, we focused on FC seeded from regions within reward circuitry (mesolimbic, mesocorticol) and within negative affect circuitry: nucleus accumbens (NAcc), hippocampus (hipp), dorsal striatum (dstri), amygdala (amyg), anterior insula (AI), dorsal anterior cingulate cortex (dACC), subgenual cingulate cortex (sgACC), posterior cingulate cortex (PCC), ventromedial prefrontal cortex (vmPFC), and thalamic subregions, including anteromedial thalamus (AM), anteroventral thalamus (AV) and mediodorsal thalamus (MD). Parcellations of the Brainnetome atlas assigned to the above regions are listed in Supplementary Table 1.

#### Statistical Analysis

Data collected at multiple times per visit, including the CADSS, VAS, and cortisol assessed via saliva samples, were analyzed using a linear mixed effects model (LMM), with dosage (placebo, 0.05 mg/kg, and 0.5 mg/kg), time (pre- and post-infusion for CADSS, BPRS; pre-, during- and post-infusion for VAS; T = −30min, 0, 30min, 60min and 150min for cortisol) and interaction of time and dosage as fixed effects. A random intercept was added for each subject to account for the repeated measures. Data collected only at the post-infusion timepoint, including the DAS, DARS, SHAPS and resting-state derived connectivity metrics, were analyzed using another LMM with only dosage as the fixed effect. A random intercept was also added for each subject to account for the repeated measures. LMM was conducted via the lmer package (https://cran.r-project.org/web/packages/lme4/index.html) in R version 4.0.5 (https://www.r-project.org/). Exploratory post-hoc paired t-tests were also conducted between each pair of sessions (0.5 mg/kg versus placebo, 0.05 mg/kg versus placebo, and 0.5 mg/kg versus 0.05 mg/kg). To explore the link between ketamine induced resting-state brain changes and perceptual, affective, and physiological changes, repeated measures correlation analyses were run using the rmcorr package (https://cran.r-project.org/package=rmcorr). Unlike conventional inter-subject correlation analysis, the repeated measures correlation analysis calculates intra-subject correlation, which, in our case, reflects how brain measures across three visits (placebo, 0.05 mg/kg, and 0.5 mg/kg) co-vary with non-brain measures, with both modalities collected at the post-infusion timepoint. For non-brain measurements collected at both post- and pre-infusion timepoints (CADSS, VAS, and cortisol), we used the post-vs. pre-infusion change for correlation analysis, denoted by Δ in figures. To retain a complete picture of ketamine-induced neuroimaging states, we did not implement corrections for multiple comparisons considering the relatively small sample size (n = 7), the pre-specified focus of our analysis strategy and because we are not intending to make inferences that generalize to a population. These interim analyses will be tested for reproducibility in a final sample of at least 20 subjects.

## Results

### Dose-dependent altered dissociative states are observed during acute ketamine administration

Using the CADSS, we observed a significant time by dose effect in dissociation (F = 7.61, p = 0.003; Fig. 2A). An increase in dissociation was reported acutely during the high dose (0.5mg/kg) condition in 4 out of 7 subjects (57%) but minimally or not at all during the low dose (0.05mg/kg) and placebo conditions. Intoxication assessed by the VAS also reflected a dose by time effect (F = 122.28, p < 0.0001; Fig. 2B). These data validate the rigor of the experimental design and confirm the placebo condition is functioning as planned. Higher dissociation correlated with higher intoxication (Repeated Measures Correlation r = 0.58, p = 0.03; Fig. 2C), suggesting partial overlap in these perceptions.

**Figure 2.**
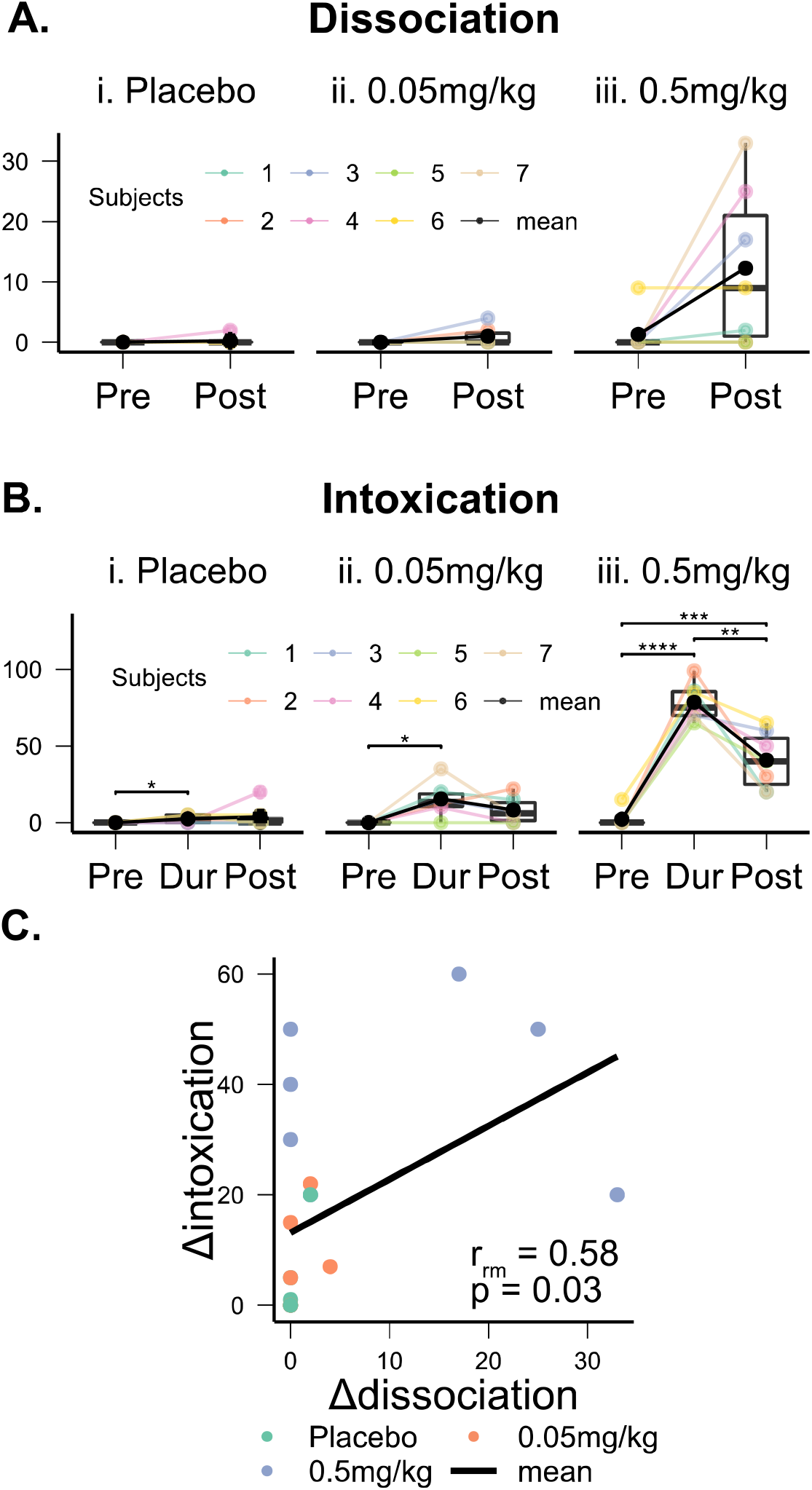
Dissociation (A) and intoxication (B) induced by ketamine. Increases induced by the 0.5mg/kg dose in CADSS-assessed dissociation were correlated with higher VAS-assessed intoxication. Color represents individual subjects, black the group mean. Abbreviations: Pre = pre-infusion; Post = post-infusion; Dur = during infusion (20-min after the start of the infusion); rm = repeated measures. *: p < 0.05, **: p < 0.01, ***: p < 0.001 as measured by a t-test.

### Dose-dependent altered affective reward responsiveness is observed during acute ketamine administration

We observed a dose-dependent effect of ketamine on reducing emotional insensitivity on the DAS (F = 8.33, p = 0.004; Fig. 3A) but not executive capacity or initiation of behaviors and thoughts. This reduction was specifically associated with intoxication but not dissociation (Repeated Measures Correlation r = −0.57, p = 0.03; Fig. 3B).

**Figure 3.**
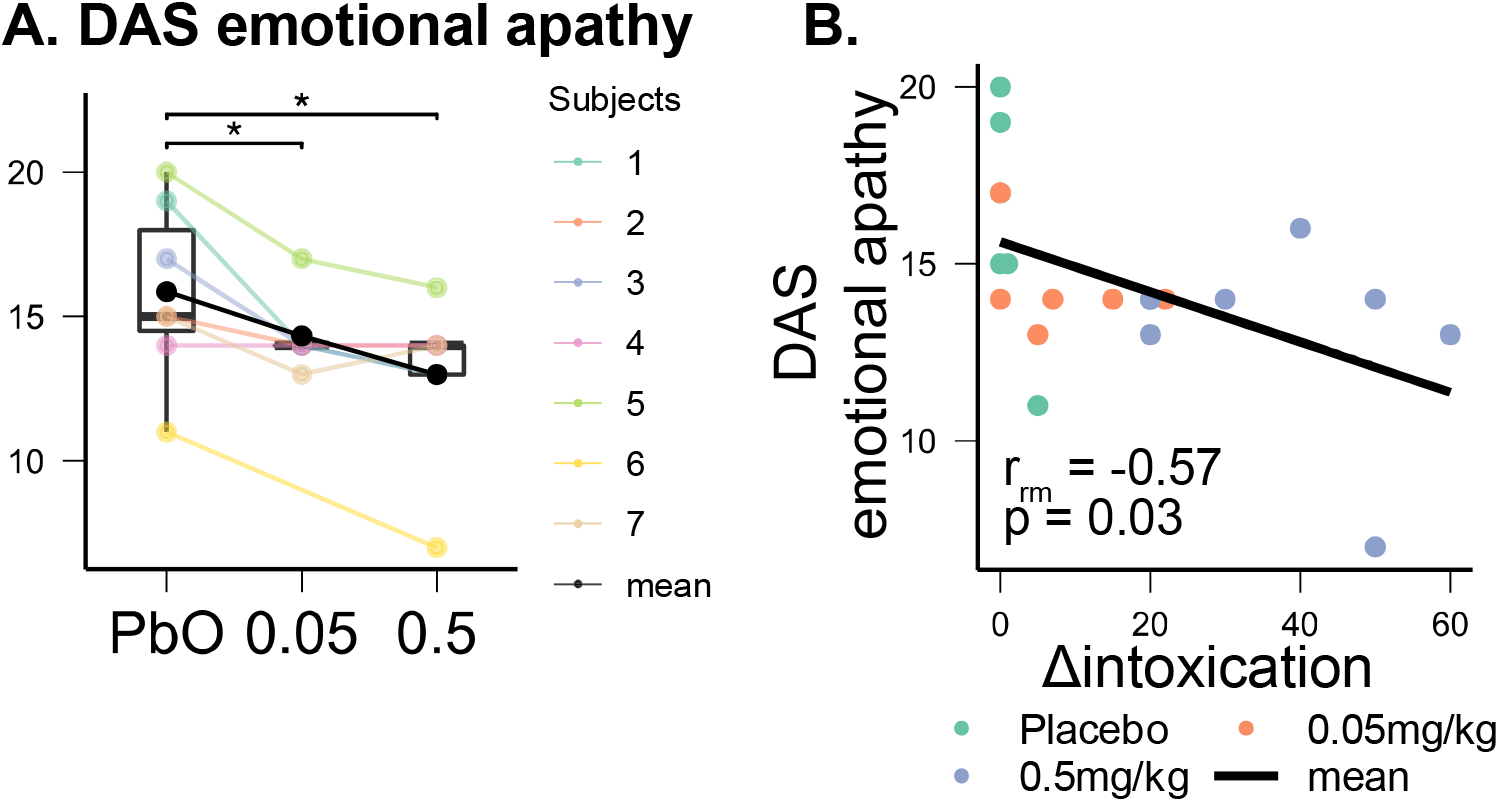
Dose-dependent ketamine-induced reductions in DAS-assessed emotional apathy (A) correlated with increased intoxication (B). Abbreviations: DAS = Dimensional Apathy Scale; PbO = Placebo; 0.05 = 0.05mg/kg; 0.5 = 0.5mg/kg; rm = repeated measures. *: p < 0.05 as measured by a paired t-test.

### Dose-dependent affective states of stress are observed during acute ketamine administration

We showed that cortisol rose by 20-30mins post-infusion and peaked by 60mins post-infusion in ketamine-induced states. This effect was pronounced in the 0.5mg/kg dose condition (F = 4.51, p < 0.001; Fig. 4A), suggesting that this condition, but not the 0.05mg/kg dose, may produce physiological stress (potentially aversive) effects. As expected, these increases in cortisol were also found to correlate with self-reported stress (DASS stress scale; Repeated Measures Correlation r = 0.67, p = 0.009; Fig. 4B). Ketamine-induced increases in cortisol were also found to correlate with greater intoxication but <u>not</u> with dissociation (Repeated Measures Correlation r = 0.81, p < 0.001; Fig. 4C).

**Figure 4.**
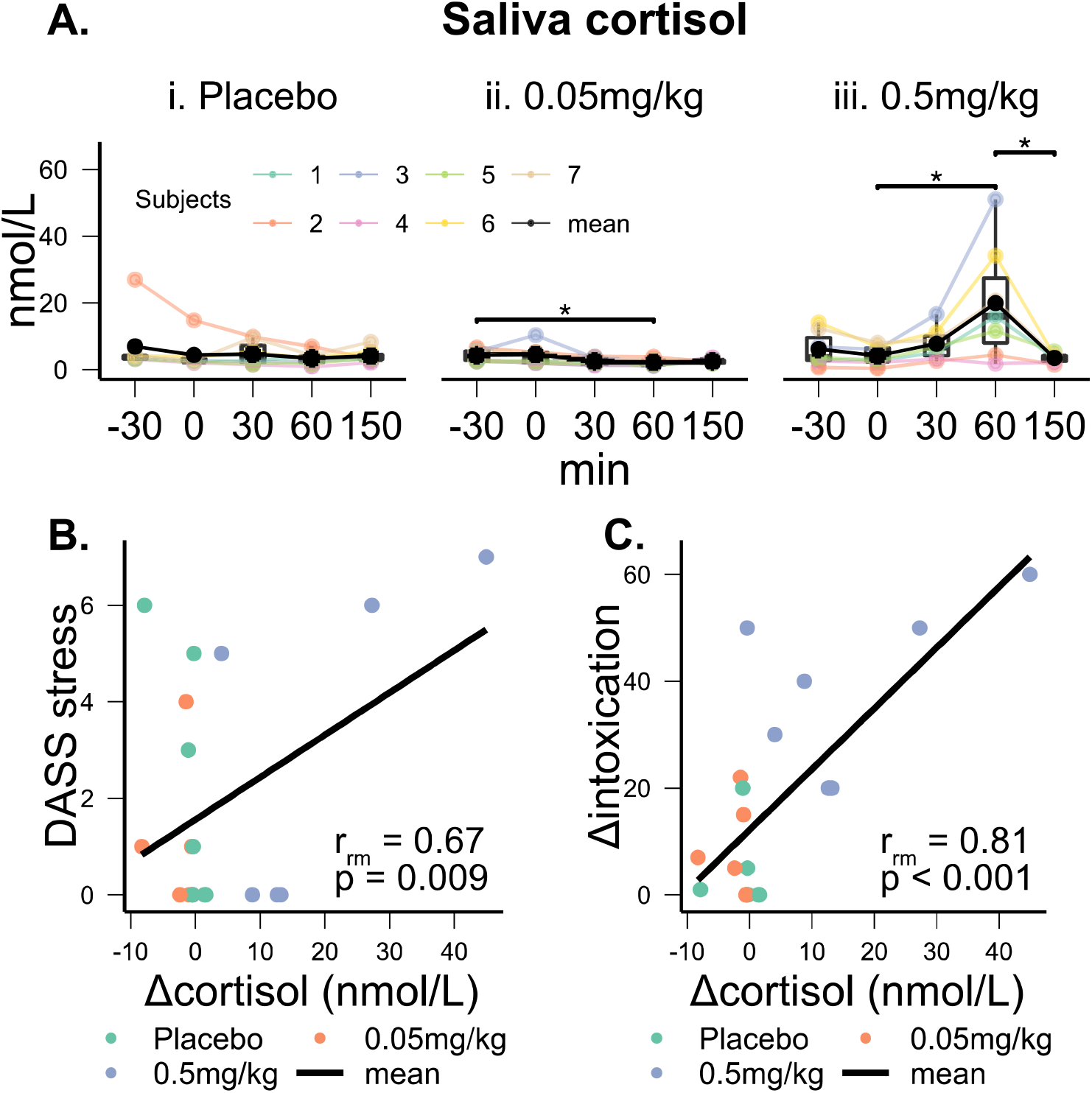
Cortisol (nmol/L) and its association with intoxication and self-reported stress under placebo and two doses of ketamine. (A) Ketamine induced dose-dependent increases in cortisol, peaking for 0.5mg/kg. X-axis labels indicate the time in minutes relative to the start of infusion (T = 0). Colors represent individual subjects, black the group mean. Ketamine-induced cortisol was significantly correlated with increased DASS-assessed stress (B) and with increased intoxication level (C). Abbreviations: DASS = Depression Anxiety Stress Scales; rm = repeated measures. *: p < 0.05 as measured by a paired t-test.

### Global functional brain connectivity altered in response to acute ketamine administration

We found a tendency toward an increase in average FC over the whole brain under the acute administration of ketamine as measured by global efficiency (Fig. 5).

**Figure 5.**
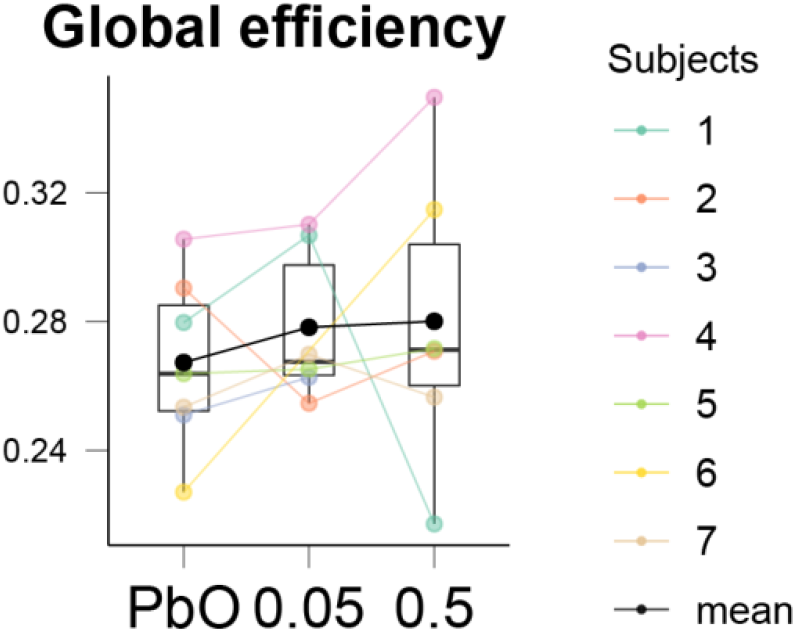
Global efficiency increased with increasing ketamine dose, on average. Colors represent individual subjects, black the group mean.

### Functional brain connectivity between thalamic sub-regions and regions of reward and negative affect circuits altered in response to acute ketamine administration

During ketamine administration, we observed drug-altered increases in functional coupling between the left anteroventral thalamus and amygdala (F = 6.91, p=0.009; Fig. 6A) and drug-altered decreases in functional coupling of the left anteroventral thalamus with the dorsal anterior cingulate cortex (F = 3.96, p = 0.046; Fig. 6B). The right thalamus, similar to the left anteroventral thalamus, showed increased coupling with the amygdala under 0.5mg/kg ketamine (F = 7.45, p = 0.007; Fig. 6C).

**Figure 6.**
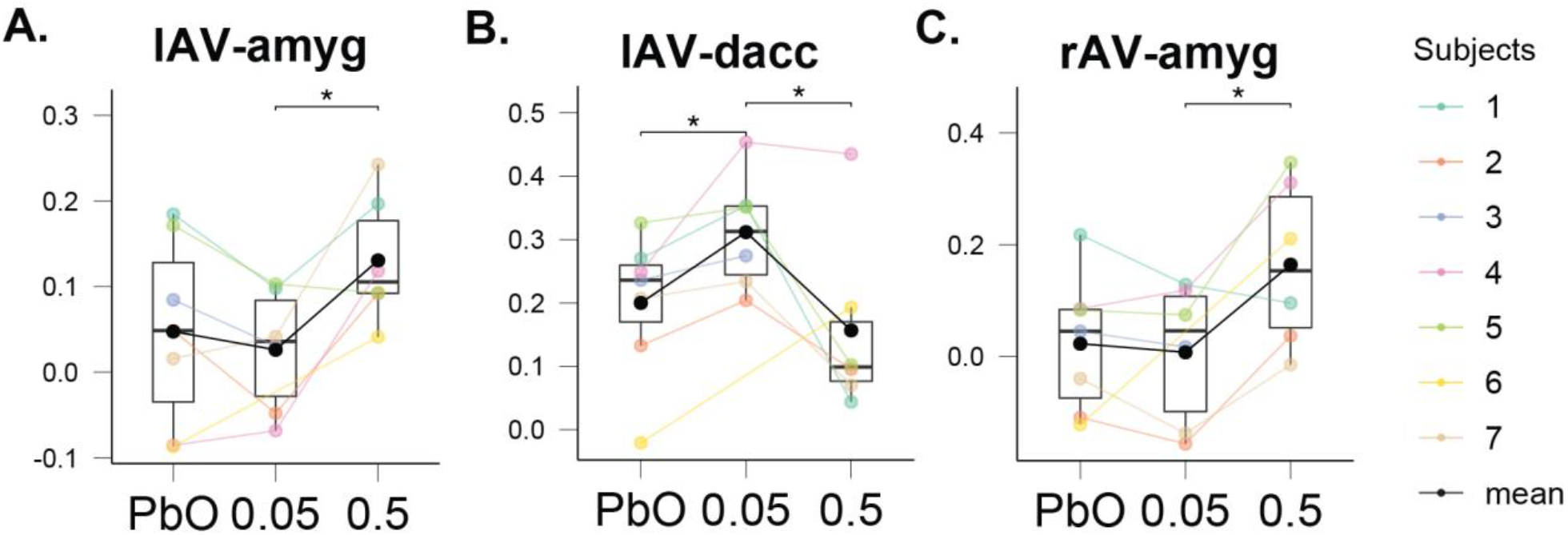
(A) Functional connectivity between the left anteroventral thalamus (lAV) and amygdala, (B) functional connectivity between the lAV and dorsal anterior cingulate, (C) functional connectivity between the right anteroventral thalamus (rAV) and amygdala. Colors represent individual subjects, black the group mean. *: p < 0.05 as determined by a t-test.

The nucleus accumbens was found to generally decrease FC with several subregions of the thalamus under ketamine conditions, including the left anterior medial thalamus (Fig. 7A), the right anterior medial thalamus (Fig. 7B), the right anteroventral thalamus (Fig. 7C), the left medial dorsal thalamus (Fig. 7D), the right medial dorsal thalamus (Fig. 7E), and the right posterior medial thalamus (Fig. 7F) (p’s > 0.05).

**Figure 7.**
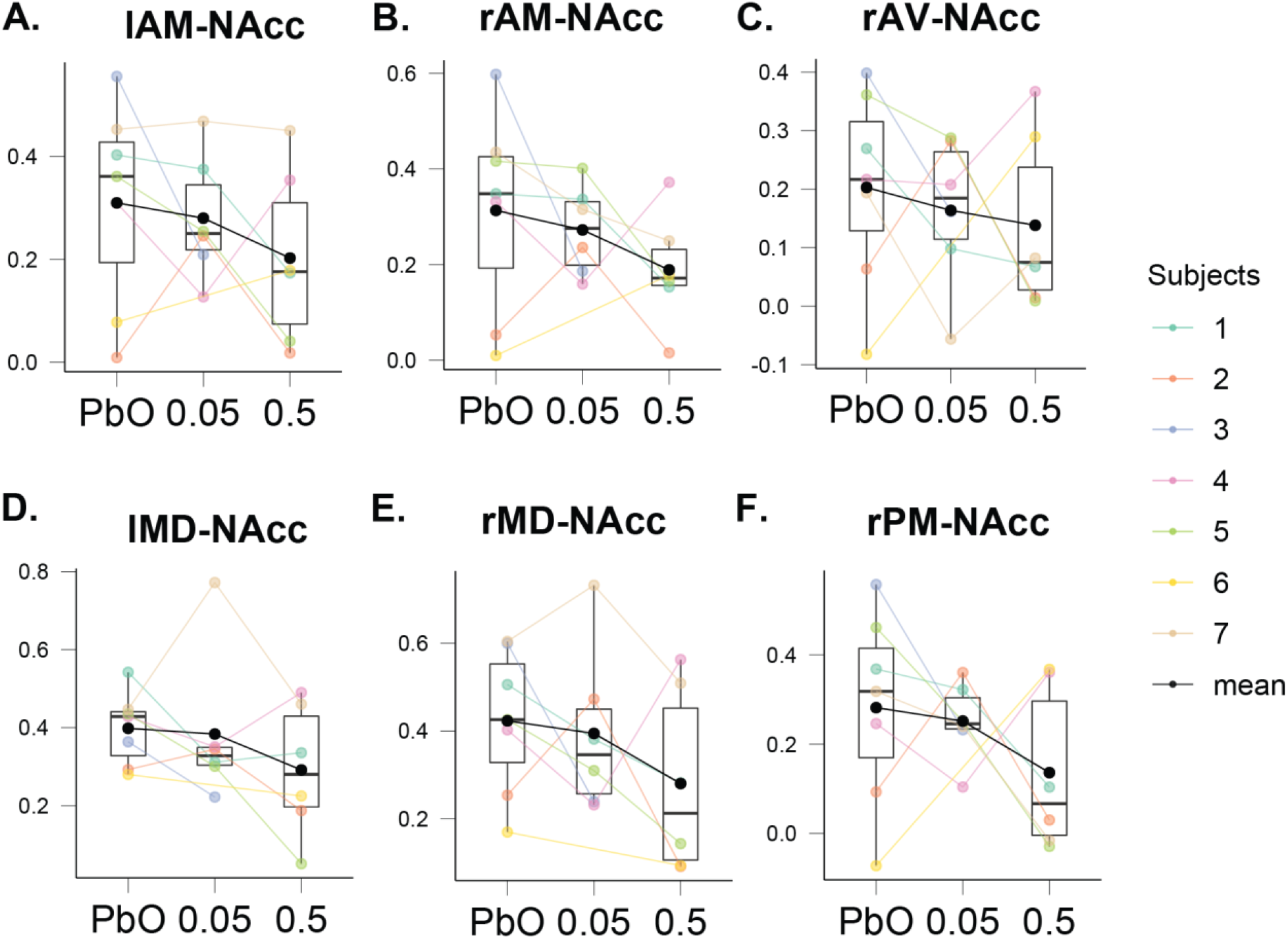
(A) Functional connectivity of the left anterior medial thalamus (lAM) and nucleus accumbens (NAcc), (B) functional connectivity of the right anterior medial thalamus (rAM) and NAcc, (C) functional connectivity of the right anteroventral thalamus (rAV) and NAcc, (D) functional connectivity of the left medial dorsal thalamus (lMD) and NAcc, (E) functional connectivity of the right medial dorsal thalamus (rMD) and NAcc, (F) functional connectivity of the right posterior medial thalamus (rPM) and NAcc. Colors represent individual subjects, black the group mean.

Additional consistent trends for ketamine-induced alternations in connectivity involving both NAcc and hippocampus regions are shown in Supplementary Results and Figures 1 and 2.

### Functional brain connectivity involving intra-thalamic sub-regions altered in response to acute ketamine administration

Thalamic sub-region to sub-region connectivity was generally characterized by a lowering of FC under ketamine conditions. This is illustrated by connections from the left anterior medial thalamus to the right anterior medial thalamus (Fig. 8A), the right anteroventral thalamus (Fig. 8B), the left and right posterior lateral ventral thalamus (Fig. 8C, 8D), the left and right posterior parietal thalamus (Fig. 8E, 8F), the left occipital posterior thalamus (Fig. 8G), and the right posterior medial ventral thalamus (Fig. 8H) (p’s > 0.05).

**Figure 8.**
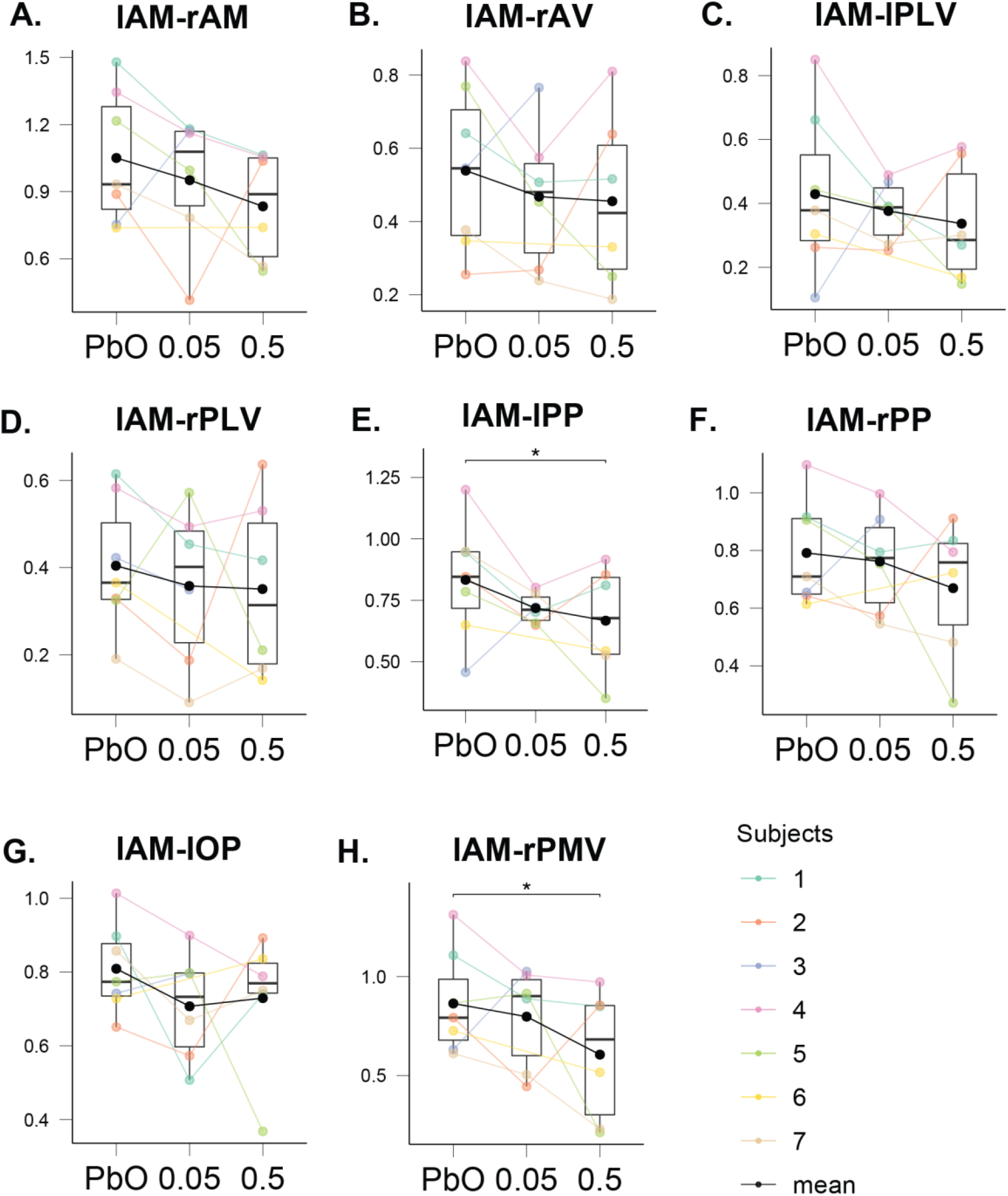
(A) Functional connectivity between the left anterior medial thalamus (lAM) and right anterior medial thalamus (rAM), (B) functional connectivity between the lAM and right anteroventral thalamus (rAV), (C) functional connectivity between the lAM and left posterior lateral ventral thalamus (lPLV), (D) functional connectivity between the lAM and right posterior lateral ventral thalamus (rPLV), (E) functional connectivity between the lAM and left posterior parietal thalamus (lPP), (F) functional connectivity between the lAM and right posterior parietal thalamus (rPP), (G) functional connectivity between the lAM and left occipital parietal thalamus (lOP), (H) functional connectivity between the lAM and right posterior medial ventral thalamus (rPMV). Colors represent individual subjects, black the group mean. *: p < 0.05 as measured by a paired t-test.

The right anterior medial thalamus also displayed a pattern of lower FC under ketamine with the right medial dorsal thalamus (Fig. 9A), the right posterior parietal thalamus (Fig. 9B), the left occipital parietal thalamus (Fig. 9C), and the right posterior medial ventral thalamus (Fig. 9D) (p’s > 0.05).

**Figure 9.**
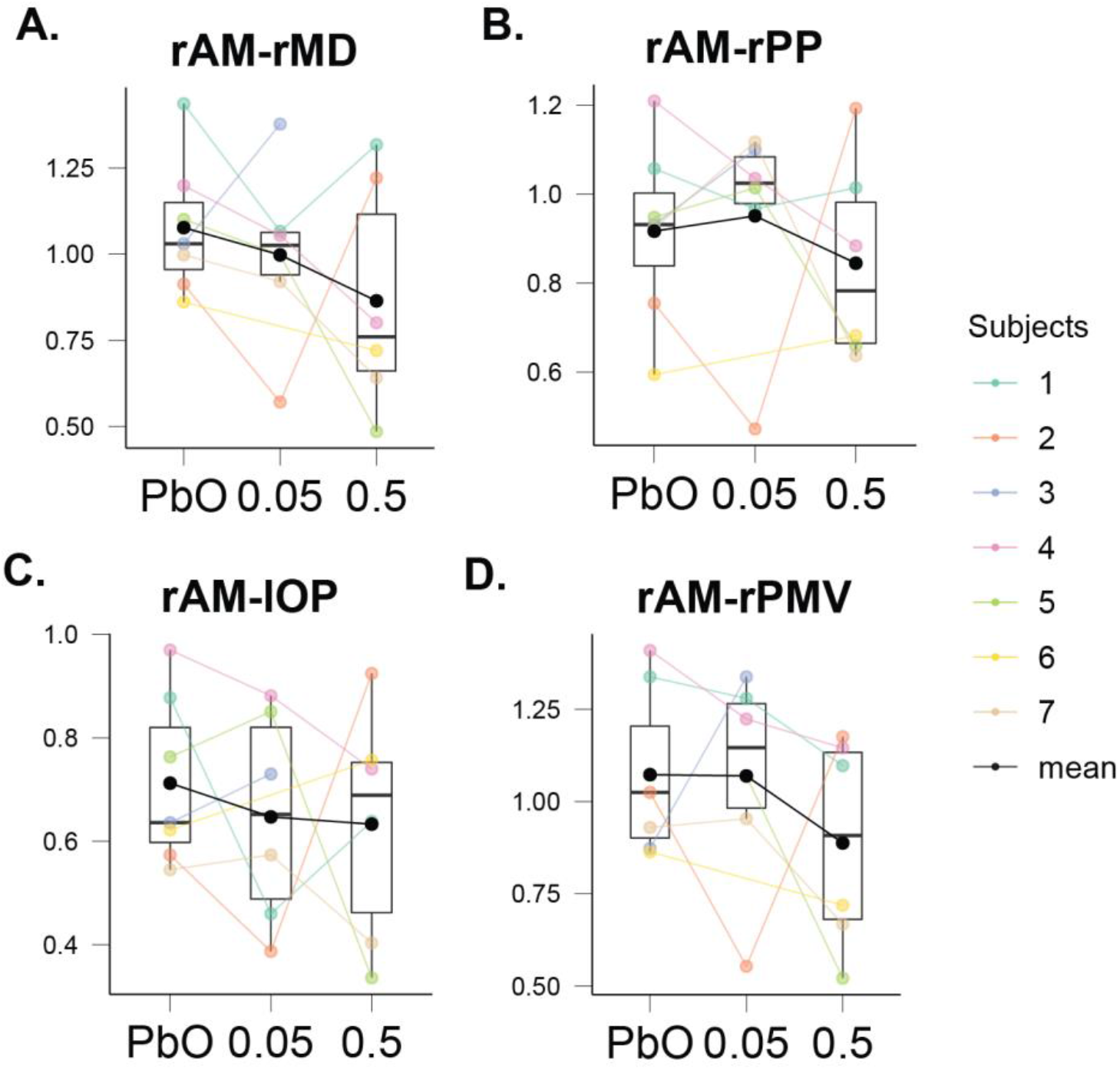
(A) Functional connectivity of the right anterior medial thalamus (rAM) and the right medial dorsal thalamus (rMD), (B) functional connectivity of the rAM and the right posterior parietal thalamus (rPP), (C) functional connectivity of the rAM and the left occipital parietal thalamus (lOP), (D) functional connectivity of the rAM and the right posterior medial ventral thalamus (rPMV). Colors represent individual subjects, black the group mean.

The left posterior medial ventral thalamus showed a distinctive although non-significant pattern of lower correlations within other regions of the thalamus including the left occipital parietal thalamus (Fig. 10A), the left posterior parietal thalamus (Fig. 10B), and the left posterior medial dorsal thalamus (Fig. 10C). The right posterior medial ventral thalamus also displayed a trend of decreased connectivity with the left occipital parietal thalamus (Fig. 10D) (p’s > 0.05).

**Figure 10.**
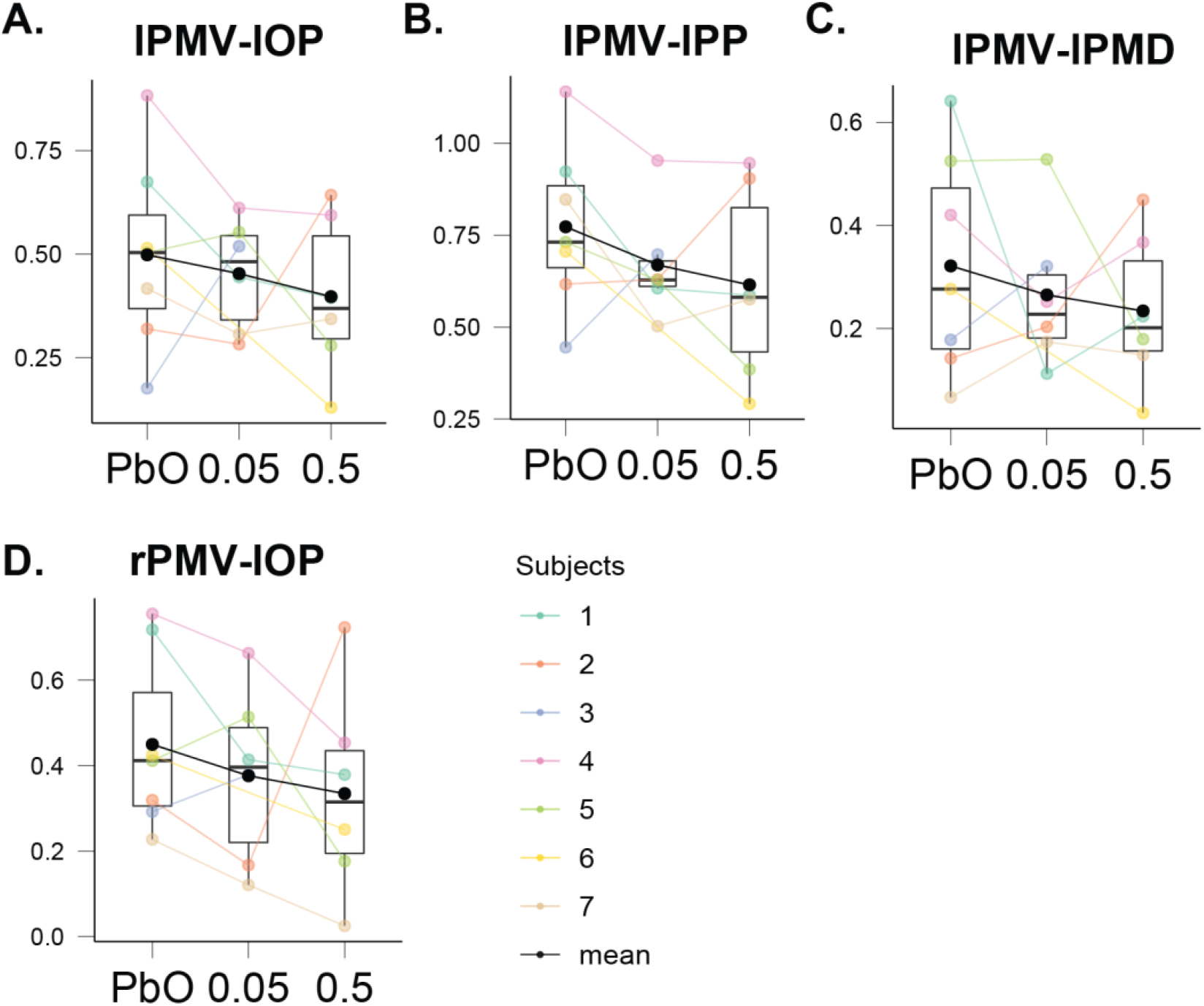
(A) Functional connectivity of the left posterior medial ventral thalamus (lPMV) and the left occipital parietal thalamus (lOP), (B) functional connectivity of the lPMV and the left posterior parietal thalamus (lPP), (C) functional connectivity of the lPMV and the left posterior medial dorsal thalamus (lPMD), (D) functional connectivity of the right posterior medial ventral thalamus (rPMV) and the left occipital parietal thalamus (lOP). Colors represent individual subjects, black the group mean.

FC changes under the influence of ketamine tended to display highly individual FC patterns, in which some subjects showed a dose-dependent increase in FC while others showed a dose-dependent decrease. This is illustrated by the connections between the left mediodorsal thalamus and right posterior mediodorsal thalamus (Fig. 11A) as well as the right mediodorsal thalamus and the right posterior thalamus (Fig. 11B) (p’s > 0.05).

**Figure 11.**
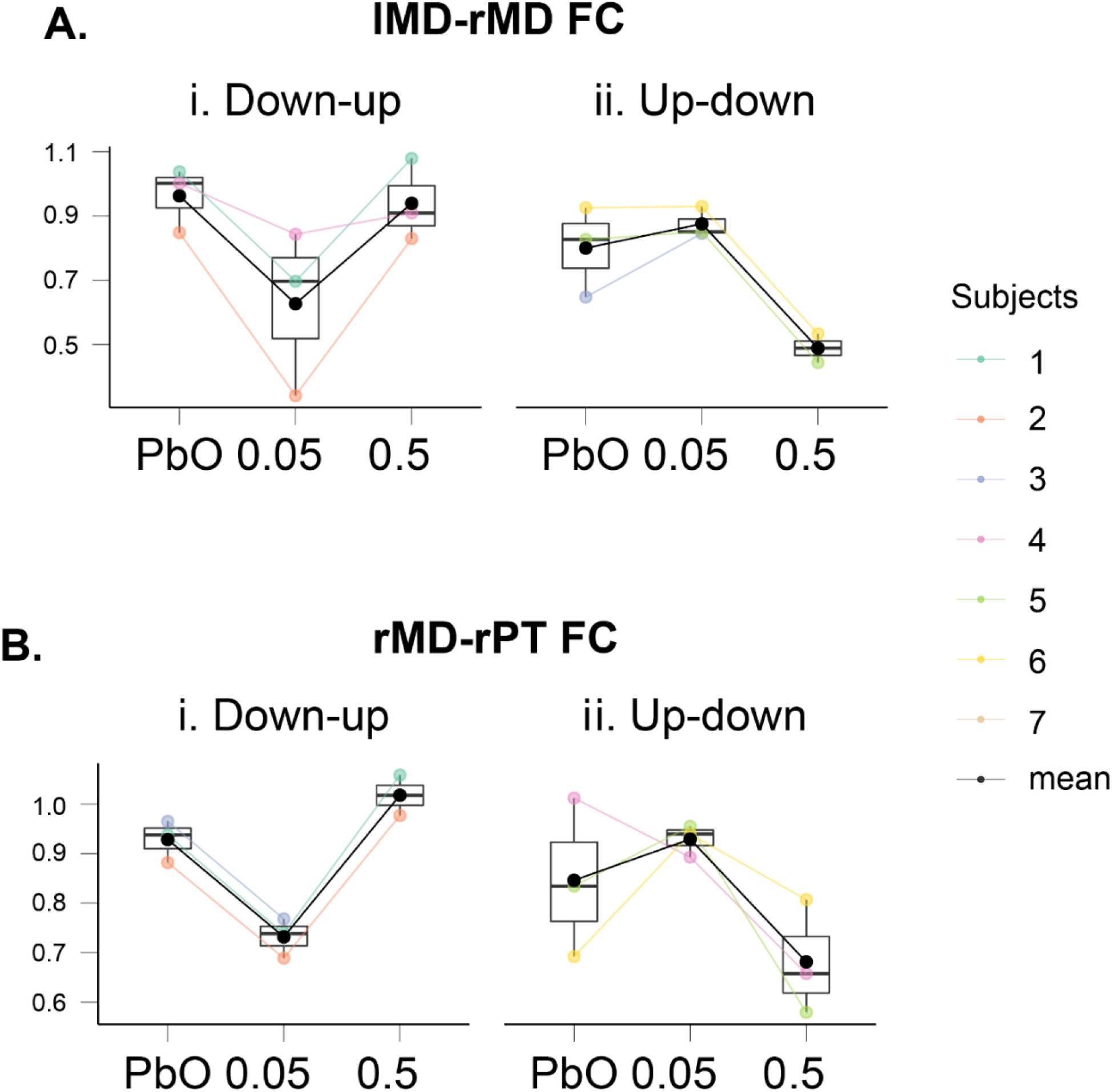
Individual variability in thalamic functional connectivity as illustrated by connections between the left mediodorsal thalamus (lMD) and right mediodorsal thalamus (rMD, A), and the rMD and right posterior thalamus (rPT, B). Colors represent individual subjects, black the group mean.

### Functional brain connectivity correlations with dissociative and affective responses

We tested whether altered states of functional brain connectivity between our regions of interest correlated with our measures of dissociative and affective responses (Fig. 12).

**Figure 12.**
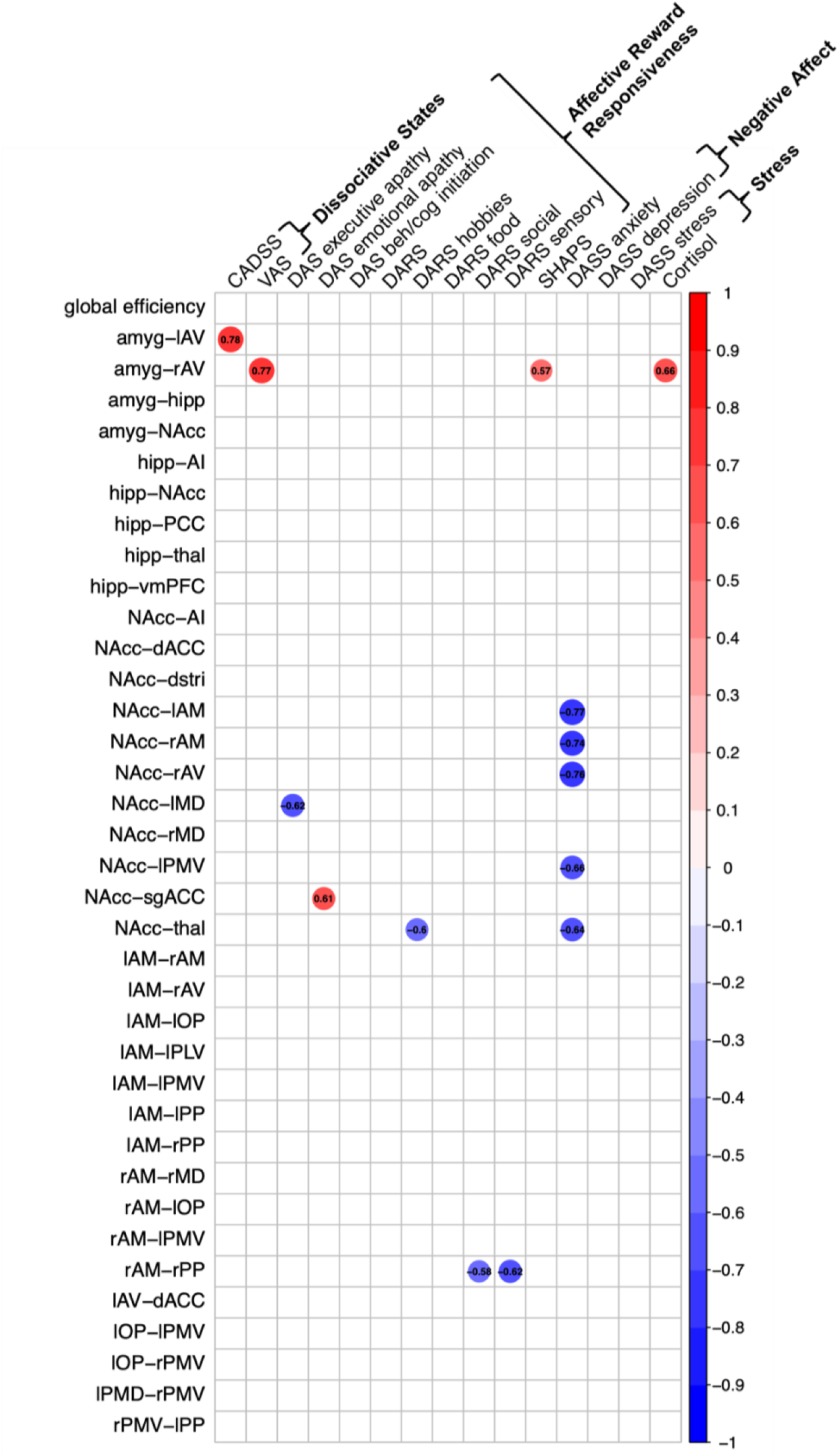
Correlations of functional connectivity measures with dissociative and affective measures. Color scale and numbers represent repeated measures r. Abbreviations: hipp = hippocampus; amyg = amygdala; AI = anterior insula; NAcc = nucleus accumbens; PCC = posterior cingulate cortex; thal =thalamus; vmPFC = ventromedial prefrontal cortex; dstri = dorsal striatum; dACC = anterior cingulate cortex; sgACC = subgenual cingulate cortex; AM = anteromedial thalamus; AV = anteroventral thalamus; MD = mediodorsal thalamus; PMV = posterior medial ventral thalamus; PLV = posterior lateral ventral thalamus; PP = posterior parietal thalamus; OP = occipital parietal thalamus; PMD = posterior medial dorsal thalamus; l = left; r = right.

#### Ketamine-altered connectivity and dissociation

Ketamine-induced increases in connectivity of the amygdala to the anteroventral sub-region of the thalamus were associated with increases in dissociation (assessed by the CADSS) and related states of intoxication (assessed by the VAS) (Fig. 12). Interestingly, and by contrast, ketamine-induced decreases in coupling of the posterior parietal thalamus and anteromedial subregion of the thalamus were associated with increased sensory altered states indicative of dissociation (assessed by the sensory domain of the DARS) (Fig. 12).

#### Ketamine-altered connectivity and affective measures related to reward responsiveness

Specific correlations were observed between affective measures relevant to reward responsiveness or its absence, and drug-altered changes in localized functional connections involving the NAcc, amygdala, and thalamic sub-regions (Fig. 12). Ketamine-induced decreases in right mediodorsal thalamus to NAcc coupling (Fig. 7E) were associated with an amelioration of executive apathy, decreases in NAcc to sgACC connectivity (Supplementary Fig. 2F) with amelioration of emotional insenstivity, and ketamine-induced increases in right anteroventral thalamus connectivity with amygdala (Fig. 6C) with increases in SHAPS-assessed pleasure. Ketamine-induced reductions in intra-thalamic connectivity between right anteromedial and left posterior parietal thalamus (Fig. 9B) were also correlated with increases in DARS-assessed sociability.

#### Ketamine-altered connectivity and affective measures related to negative affect

We discovered a consistent profile of associations between ketamine altered connectivity involving NAcc, amygdala, dACC and specific thalamic sub-regions and effects of anxiety (Fig. 12). Ketamine-altered increases in coupling of the right anteroventral thalamus and amygdala (Fig. 12) were associated with increases in cortisol, an indicator of biochemical stress (Fig. 12).

## Discussion

In this report, we present preliminary results of brain altered states induced acutely by two doses of subanesthetic IV ketamine and the dissociative and affective responses that accompany these altered states. Deconstructing these brain-response states at both the group and individual subject level is essential to understanding both the abuse liability of ketamine and its therapeutic potential.

Our first set of analyses revealed robust, dose-dependent effects of acute ketamine administration on dissociative and affective responses. Ketamine induced a clear increase in dissociation and subject-reported intoxication. The dose-dependent nature of this dissociative effect, in which dissociation peaked during the higher dose ketamine (0.5 mg/kg), was present but to a lesser extent during the lower dose (0.05 mg/kg), and was not present during placebo demonstrates the rigor of our scientific design and placebo control. Dissociation was also highly correlated with intoxication, consistent with the dose-dependent effects of ketamine on dissociation. Regarding affective reward-related responses, there was a dose-dependent effect of ketamine on ameliorating emotional insensitivity, which was associated with changes in intoxication but not dissociation. For affective stress responses, we observed a striking dose-dependent increase in cortisol, peaking at 60 minutes post-infusion and correlated with self-reported stress and greater intoxication, but not with dissociation.

Analyses of ketamine-altered states of functional brain connectivity revealed a tendency towards increasing brain-wide global efficiency, which suggests that ketamine may increase connections between long-range nodes that support many other connections and help to minimize inefficient localized connections. Indeed, we observed a consistent effect of ketamine on reducing localized functional connections involving the nucleus accumbens and the prefrontal regions that define the reward circuit. Despite the small group of subjects in this preliminary investigation, the general consistency of these findings overall as well as in individuals suggests that these patterns may extend to larger samples. Previous studies investigating FC during ketamine administration have also reported reduced FC in the reward circuit [18, 54, 55]. These effects may reflect the impact of ketamine on glutamate transmission given connectivity of regions within this circuit has been shown to be modulated by changes in glutamatergic neurotransmission [24].

Our analyses of correlations between dissociative/affective responses and ketamine-altered brain connectivity states suggested that these responses can be deconstructed into specific brain connectivity-response signatures. For example, on the one hand, ketamine-induced dissociation was correlated specifically with FC of the amygdala to the left anteroventral thalamus. Affective states of stress, on the other hand, were correlated with ketamine-induced coupling of the amygdala to the right anteroventral thalamus. Dissociation and cortisol changes were themselves uncorrelated. Thus, specific signatures of localized coupling, possibly lateralized, may underlie distinct states of dissociation versus stress. Regarding reward-related affective responses, FC of the nucleus accumbens to the subgenual anterior cingulate cortex was correlated with a reduction in emotional insensitivity, consistent with previous reports implicating reward circuitry (including connections between the nucleus accumbens and cingulate regions of the medial prefrontal cortex) in the modulation of emotional and motivational states [56].

To advance our understanding of these complex states and drug-related brain mechanisms, it is increasingly important in both basic and clinical neuroscience to quantify inter- and intra-subject variability. In doing so, the opportunity to directly translate across rodent models and human subjects findings is also enhanced. Our findings suggest that some measures, such as cortisol increases, reveal relatively consistent ketamine-induced effects. These measures offer a means to anchor the interpretation of effects that may vary more between and within subjects. At baseline, we observe that FC for some connections of interest formed at least two types of subject signatures. Thus, taking account of variability at baseline should be one focus of future acute ketamine studies.

Our findings may have implications for the potential therapeutic use of ketamine. There are mixed findings and views as to whether dissociation is necessary to achieve antidepressant effects of ketamine [7, 8] or whether these effects are considered as adverse events. An answer may lie in the personalized responses of each individual person, which might require in-depth analysis of individual variation. Relatedly, it is also unclear whether dissociative effects drive the addictive potential of ketamine, and if so, why. Notably, ketamine administration elicited large increases in cortisol in our healthy subjects, unlike cortisol reductions observed in depressed individuals, who might already enter similar studies with elevated cortisol. Thus, anchoring analyses by baseline cortisol may help further deconstruct this individual variation. Given the consistent associations observed between dose-dependent ketamine-altered brain connectivity and anxiety, future studies may seek to stratify healthy subjects by anxiety and, in treatment studies of clinical subjects, consider focusing on the lower dose of 0.05 mg/kg to minimize anxiety in these subjects [6]. Our focus on reward and negative affect circuitry may also have implications for potential therapeutic applications of ketamine, since disruptions in these circuits have been reported in clinical depression and anxiety disorders, as well as accompanying substance use dependence [55]. As the number of subjects in this study increases, a key next step will involve exploring whether variability in baseline affective states (including self-reported anxiety and cortisol-assessed stress) and functional brain connectivity contribute to subsequent changes in these responses and connectivity signatures as a function of acute ketamine administration.

The current study has several limitations. First, because this is a preliminary report, the sample size is relatively small. The current sample was not intended to provide a basis for making broader inferences. We will determine the reproducibility and effect size of results within stringent confidence intervals as we complete the final phase of this study. A related limitation involves the limited range of scores on some of the affective response measures of interest. While adhering to the inclusion and exclusion criteria, we seek to expand the range of scores as the sample increases in the final study phase. While we have made every effort to administer the questionnaires and conduct scanning at the same time points for each individual, a final limitation is that this was not always possible due to needing to first address adverse events or other issues for some participants (such as nausea or scanner delays).

In summary, this study is the first to provide an in-depth characterization of the relationships between key dissociative and affective responses to acute ketamine administration at different doses in healthy subjects and to chart neural correlates of these responses. Understanding the neural mechanisms underlying the acute response to ketamine is important for the safe therapeutic use of ketamine given the relationship of these effects with both its abuse liability and treatment efficacy.

## Acknowledgements

The authors would like to extend their appreciation to the participants in this study. We thank Dr. Robert Welsh for the use of the ConnTool Functional Connectivity Toolbox.

## Clinicaltrials.gov

NCT03475277

## Ethics Approval and Consent to Participate

The Institutional Review Board of Stanford University has approved this protocol (#41173). A study coordinator thoroughly explained the protocol to participants and answered any questions before they provided written informed consent to begin the study. The study was conducted according to the principles of the Declaration of Helsinki.

## Authors’ Contribution

CIR, BK, and LMW developed the study; XZ and KGW performed the data analysis. LMH, XZ, KGW, and LMW wrote the manuscript, and all authors reviewed and approved the manuscript.

## Data Availability Statement

Data collection is ongoing. Data will be made publicly available once data collection has been completed.

