## Supplemental Files for "Acute Ketamine Modulated Functional Brain Coupling and Dissociative and Affective States in Human Subjects: Interim Analyses"

Supplementary Results, Table, and Figures

Supplementary Table 1. Regions of Interest

| BNT Regions | ROI name | ROI number |
|---|---|---|
| Amygdala | Amyg_L_2_1 | 211 |
| Amygdala | Amyg_R_2_1 | 212 |
| Amygdala | Amyg_L_2_2 | 213 |
| Amygdala | Amyg_R_2_2 | 214 |
| Anterior Insula | INS_L_6_1 | 163 |
| Anterior Insula | INS_R_6_1 | 164 |
| Anterior Insula | INS_L_6_5 | 171 |
| Anterior Insula | INS_R_6_5 | 172 |
| dACC | CG_L_7_3 | 179 |
| dACC | CG_R_7_3 | 180 |
| dACC | CG_L_7_5 | 183 |
| dACC | CG_R_7_5 | 184 |
| dorsal striatum | BG_L_6_1 | 219 |
| dorsal striatum | BG_R_6_1 | 220 |
| dorsal striatum | BG_L_6_4 | 225 |
| dorsal striatum | BG_R_6_4 | 226 |
| dorsal striatum | BG_L_6_5 | 227 |
| dorsal striatum | BG_R_6_5 | 228 |
| dorsal striatum | BG_L_6_6 | 229 |
| dorsal striatum | BG_R_6_6 | 230 |
| Hippocampus | Hipp_L_2_1 | 215 |
| Hippocampus | Hipp_R_2_1 | 216 |
| Hippocampus | Hipp_L_2_2 | 217 |
| Hippocampus | Hipp_R_2_2 | 218 |
| NAcc | BG_L_6_3 | 223 |
| NAcc | BG_R_6_3 | 224 |
| PCC | CG_L_7_1 | 175 |
| PCC | CG_R_7_1 | 176 |
| PCC | CG_L_7_4 | 181 |
| PCC | CG_R_7_4 | 182 |
| PCC | CG_L_7_6 | 185 |
| PCC | CG_R_7_6 | 186 |
| sgACC | CG_L_7_2 | 177 |
| sgACC | CG_R_7_2 | 178 |
| sgACC | CG_L_7_7 | 187 |
| sgACC | CG_R_7_7 | 188 |
| Thalamus | Tha_L_8_1 | 231 |
| Thalamus | Tha_R_8_1 | 232 |
| Thalamus | Tha_L_8_2 | 233 |
| Thalamus | Tha_R_8_2 | 234 |
| Thalamus | Tha_L_8_3 | 235 |
| Thalamus | Tha_R_8_3 | 236 |
| Thalamus | Tha_L_8_4 | 237 |

Abbreviations: BNT = Brainnetome; dACC = dorsal anterior cingulate cortex; NAcc = nucleus accumbens; PCC = posterior cingulate cortex; sgACC = subgenual anterior cingulate cortex.

We show additional trends that involve consistent regional connections during ketamine administration and that warrant further investigation in a larger sample. These trends were localized to ketamine-induced altered connectivity involving the hippocampus and the NAcc.

During ketamine administration, the hippocampus showed a non-significant tendency to decrease in FC with multiple regions, including the amygdala, anterior insula, and NAcc, as well as a non-significant increases in FC to the posterior cingulate cortex, thalamus, and ventromedial prefrontal cortex (Supplementary Fig. 1). The NAcc generally decreased FC under ketamine conditions (not significant), including connections to the amygdala, anterior insula, dorsal striatum, dorsal anterior cingulate cortex, thalamus, and subgenual anterior cingulate cortex (Supplementary Fig. 2).

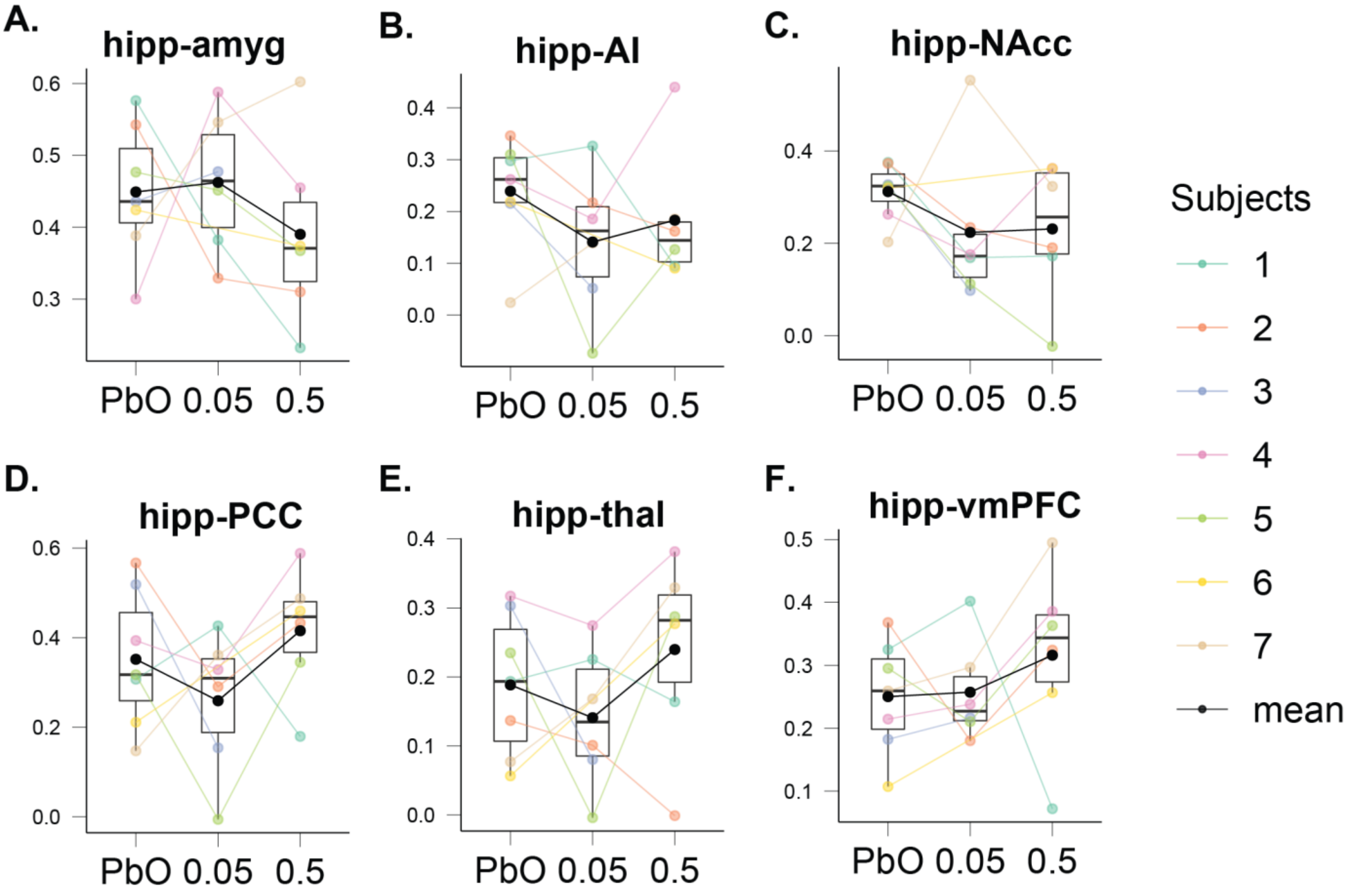


**Supplementary Figure 1**. (A) Functional connectivity between the hippocampus and amygdala, (B) functional connectivity between the hippocampus and anterior insula (AI), (C) functional connectivity between the hippocampus and nucleus accumbens (NAcc), (D) functional connectivity between the hippocampus and posterior cingulate cortex (PCC), (E) functional connectivity between the hippocampus and thalamus, (F) functional connectivity between the hippocampus and ventromedial prefrontal cortex (vmPFC). Colors represent individual subjects, black the group mean.

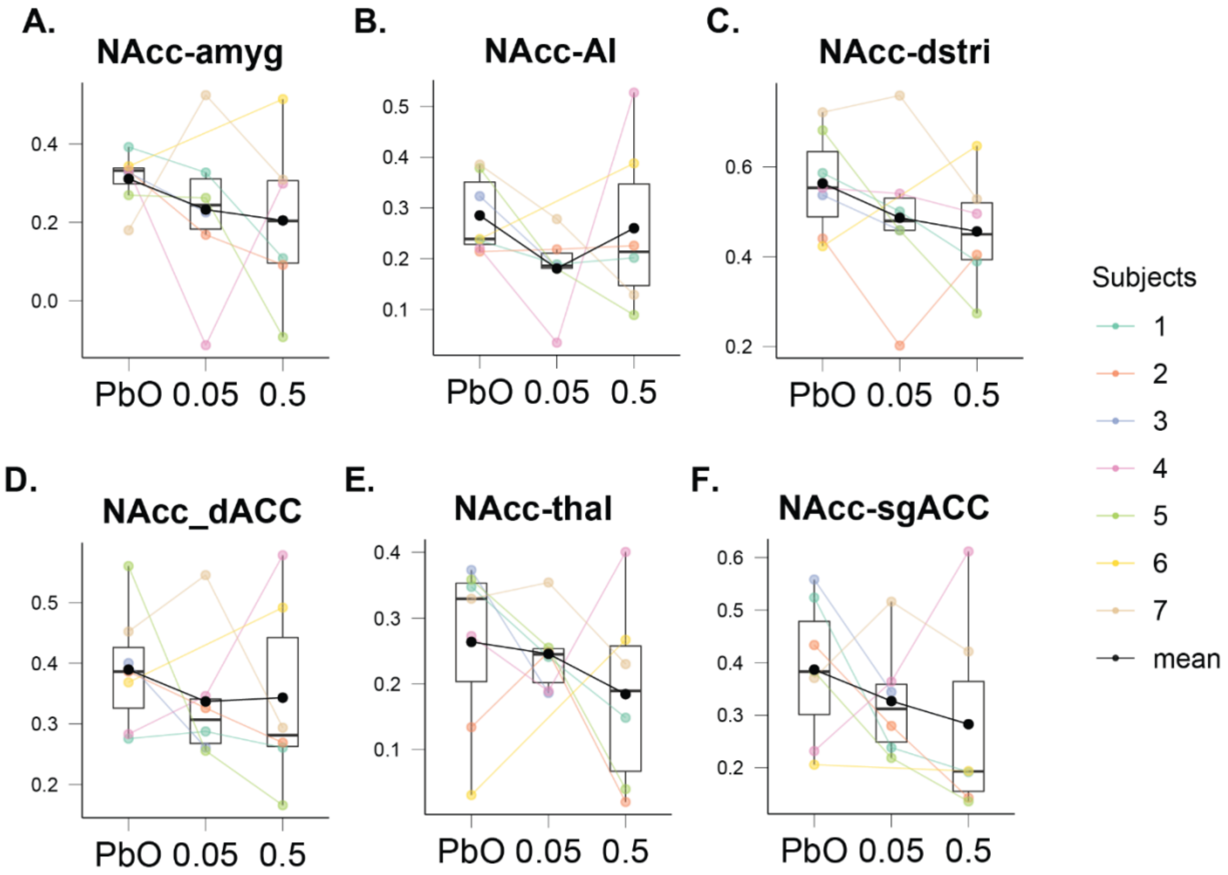


**Supplementary Figure 2**. (A) Functional connectivity between nucleus accumbens (NAcc) and amygdala, (B) functional connectivity between NAcc and anterior insula (AI), (C) functional connectivity between NAcc and dorsal striatum (dstri), (D) functional connectivity between NAcc and dorsal anterior cingulate cortex (dACC), (E) functional connectivity between NAcc and thalamus, (F) functional connectivity between NAcc and subgenual anterior cingulate cortex (sgACC). Colors represent individual subjects, black the group mean.
